# Thermostable designed ankyrin repeat proteins (DARPins) as building blocks for innovative drugs

**DOI:** 10.1101/2021.04.27.441521

**Authors:** Johannes Schilling, Christian Jost, Ioana Mariuca Ilie, Joachim Schnabl, Oralea Buechi, Rohan S. Eapen, Rafaela Truffer, Amedeo Caflisch, Patrik Forrer

## Abstract

Designed Ankyrin Repeat Proteins (DARPins) are a class of antibody mimetics with a high and mostly unexplored potential in drug development. They are clinically validated and thus represent a true alternative to classical immunoglobulin formats. In contrast to immunoglobulins, they are built from solenoid protein domains comprising an N-terminal capping repeat, one or more internal repeats and a C-terminal capping repeat. By using *in silico* analysis and a rationally guided Ala-Scan, we identified position 17 of the N-terminal capping repeat to play a key role for the overall protein thermostability. The melting temperature of a DARPin domain with a single full-consensus internal repeat was increased by about 8°C to 10°C when the original Asp17 was replaced by Leu, Val, Ile, Met, Ala or Thr, as shown by high-temperature unfolding experiments at equilibrium. We then transferred the Asp17Leu mutation to various backgrounds, including different N- and C-terminal capping repeats and clinically validated DARPin domains, such as the VEGF-binding ankyrin repeat domain of abicipar pegol. In all cases, the proteins remained monomeric and showed improvements in the thermostability of about 8°C to 16°C. Thus, the replacement of Asp17 seems to be generically applicable to this drug class. Molecular dynamics simulations show that the Asp17Leu mutation reduces electrostatic repulsion and improves van-der-Waals packing, rendering the DARPin domain less flexible and more stable. Interestingly, such a beneficial Asp17Leu mutation is present in the N-terminal caps of three of the five DARPin domains of ensovibep, a SARS-CoV-2 entry inhibitor currently in clinical development. This mutation is likely responsible, at least in part, for the very high melting temperature (>90°C) of this promising anti-Covid-19 drug. Overall, such N-terminal capping repeats with increased thermostability seem to be beneficial for the development of innovative drugs based on DARPins.

## Introduction

DARPins are a class of antibody mimetics, that have been conceived and developed about two decades ago at the University of Zurich ^1–3^. Their application as research tool and protein therapeutic was recently reviewed ^4^. Originally devised as an alternative to immunoglobulins (“antibodies”), the potential of DARPins in protein engineering, directed evolution of binders and drug development became obvious immediately at inception. Importantly, this potential extends beyond areas of applications that have classically been “occupied” by recombinant immunoglobulins. The DARPin scaffold was shown to serve as an alternative ^5^, as a complementation ^6^ and as an expansion of what is possible with binders derived from immunoglobulins ^7,8^. Translation of academic research in DARPin technology towards pharmaceutical benefits has been predominantly steered by Molecular Partners, who provided the fundamental clinical validation of the scaffold ^9^. However, in light of the long generation cycles in drug development – especially in the case of biologics that typically require ten years from concept to drug approval - the DARPin technology can still be regarded as young and emerging and the full potential of DARPins as a class of biologics has yet to be realized. The recent development of ensovibep ^10^, a multi-specific anti-SARS-CoV-2 DARPin, which has entered clinical trials in November 2020 in less than nine months after initial research and development activities had commenced, reinforces this high potential ^11,12^.

DARPins are based on natural ankyrin repeat proteins ^13^, that have evolved to mediate various kinds of protein-protein interactions in all kingdoms of life ^14^. Their structure is simpler than that of immunoglobulins. Whereas immunoglobulins naturally consist of four polypeptide chains, and unite more than four chains in recombinant formats like T-cell bispecifics (TCBs) ^15^, a single polypeptide chain is sufficient to form a multispecific DARPin ^16^. For example, ensovibep ^10^ combines five DARPin domains on a single polypeptide chain, in which two domains bind human serum albumin (HSA) and three domains associate with the SARS-CoV-2 spike protein ^11^. DARPins are built from solenoid protein domains, which possess a modular architecture that was derived by a consensus design approach ^2,17,18^: a stack of internal ankyrin repeats, each composed of 33 amino acids, flanked by N- and C-terminal capping repeats (N- and C-Caps) that function to seal the hydrophobic core of the protein domain (**Figure 1**). Together these structural units form an elongated ankyrin repeat domain. Amino acids present at defined positions at the surface of the internal repeats form a paratope, enabling the binding to target proteins with high affinity and specificity ^3,19,20^. These positions are randomized in DARPin libraries, which are used as starting point for *in vitro* selection, most prominently by means of ribosome display ^21^, to generate highly-specific target binding molecules. Originally, the N- and C-Caps of DARPins were taken from the human guanine-adenine-binding protein (hGABP_beta1) as they could be adapted to fit to the consensus designed internal repeats ^2^. Such original N- and C-Caps are also present in the first DARPin that became clinically validated - abicipar pegol. Despite the clinical validation, most of the amino acid sequence of these original caps was not optimized, indicating that there may be room for further improvements, in particular for those that increase DARPin thermostability.

**Figure 1.**
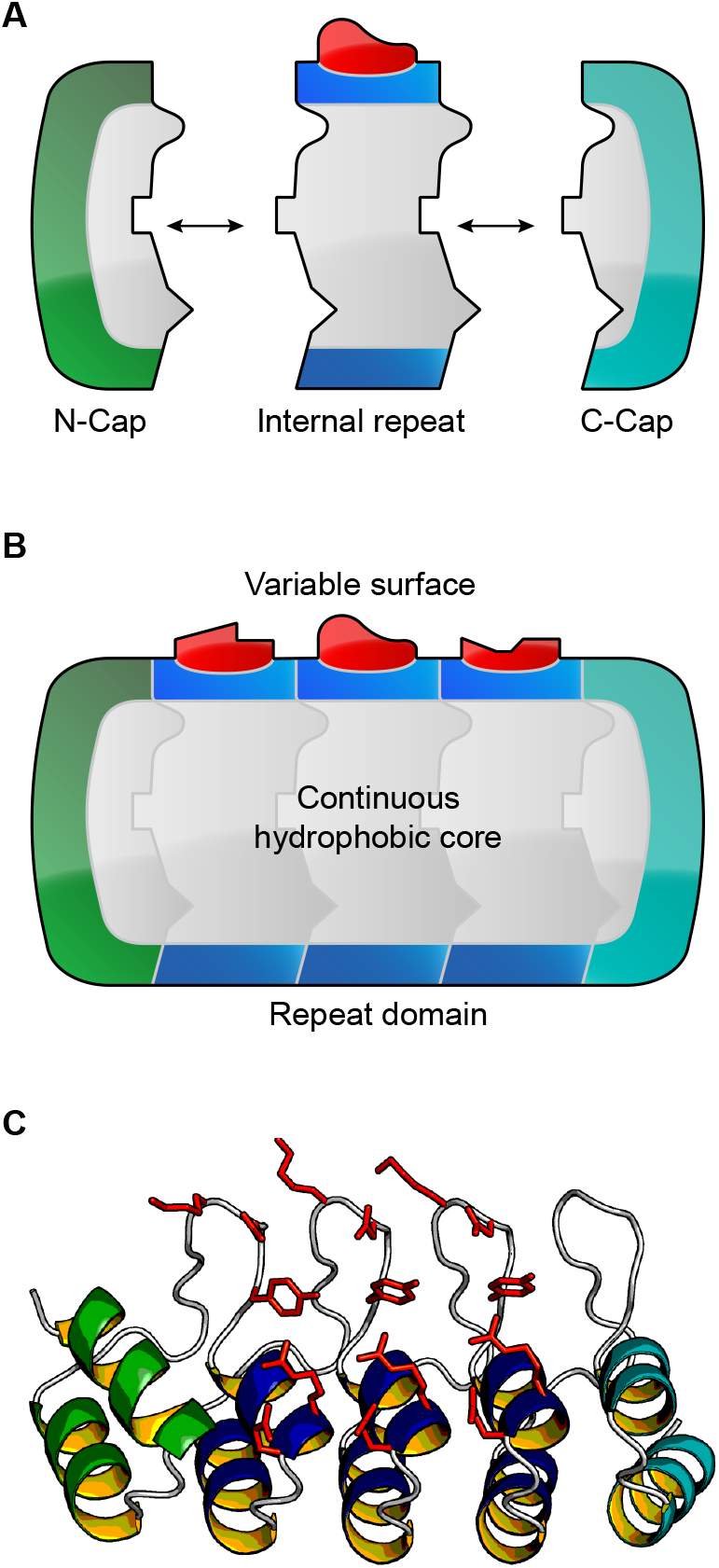
**A)** Schematic representation of a DARPin domain. Designed N- and C-terminal capping repeats and designed self-compatible repeat modules are the building blocks that stack via compatible interfaces. **B)** The repeat modules stack together forming a continuous hydrophobic core, which is sealed by an N-Cap (green) and a C-Cap (cyan). The repeat domains display variable molecular surfaces (randomized positions, depicted in red), which are potential target binding sites. ^*1,2*^ **C)** Structure of a DARPin (PDB ID: 2XEE ^*25*^) in ribbon representation with the same color scheme as above: Helices of the N-terminal capping repeat, the internal repeat modules and the C-terminal capping repeat are colored green, blue and cyan, respectively. Side chains of randomized positions are displayed as red sticks. This figure was created with open-source PyMOL ^26^.

In general, we identify three major motivations that fuel the quest for increased thermostability of biologics: (i) Reduction of aggregation and thus reduction of immunogenicity risk, (ii) simplification of chemistry, manufacturing, and controls (CMC) processes, thus bringing down the manufacturing costs, and reduction of cold-chain-drug storage requirements and (iii) increase of degrees of freedom for protein engineering to allow for mode-of-action design, for example, advanced multispecificity ^11^, receptor fine-tuning ^22^ and proximity-based activation ^23^. For the thermostability of DARPins, the importance of the capping repeats – particularly the C-Cap – was first shown by Interlandi and coworkers ^24^. Seven point mutations, five of which are located in the interface to the preceding internal repeat and two at the very C-terminus, were introduced to optimize the C-Cap and were shown to increase the melting temperature (*T*_m_) of a model DARPin, consisting of an N-Cap, a single full-consensus repeat and a C-Cap by about 17°C; *i*.*e*. from 60°C (wt) to 77°C (the respective C-Cap variant was referred to as the “mut5” C-Cap) ^24^. When this improved mut5 C-Cap was compared to the original C-Cap ^2^, a rigid-body movement of the C-Cap towards the internal repeat was observed, as evidenced by crystallographic data ^25^. This movement results in an increased buried surface area and a superior complementarity of the interface between the internal repeat and the C-Cap, which explains the improved thermostability.

While the C-Cap of DARPins was thoroughly investigated, we are not aware of any scientific article describing a corresponding analysis or thermostability improvement of the original N-Cap (denoted as “N01” N-Cap in the following, (**Figure 2**)), which is still predominantly used by the research community. Nevertheless, thermostability improvements of the N-Cap have been published in the patent application WO2012/069655 (WO’655) and are clinically validated in the aHSA domains of ensovibep ^10^. The corresponding N-Cap of WO’655 is denoted “N02” in the following (**Figure 2**). Comparing N02 with the original N-Cap, N01, resulted in a *T*_m_ increase of approximately 7°C. It is likely that this improvement mainly arises from the Met24Leu-mutation present in N02, which removes the only methionine (besides the methionine encoded by the start codon) from the original DARPin sequence, and thereby also removes this hotspot for oxidation. The N-Cap analysis of WO’655 was limited to the RILMAN sequence motif around Met24 of N01.

Here, we set out to improve the thermostability of DARPins through engineering the N-Cap. Through *in silico* analyses and high-temperature unfolding experiments at equilibrium we identify N-Cap Asp17 as an Achilles heel of DARPin domains. Molecular dynamics (MD) simulations and MD trajectory analysis provide an explanation for the significantly increased *T*_m_ values observed upon Asp17 replacement in DARPin domains.

**Figure 2.**
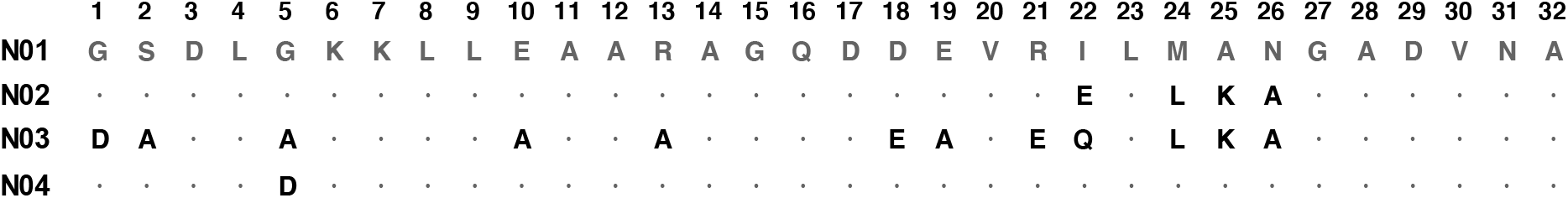
Amino acid sequence alignment of different DARPin N-Caps. Amino acids in N02, N03 and N04 that are identical to the one present in N01 are indicated as dots. The sequence numbering is shown as used in the text. N01, the original N-Cap as described by Binz et al. ^2^. N02, an improved N-Cap as described in WO’655 and present in the two anti-HSA DARPin domains of ensovibep ^10^. N03, an N-Cap where 12 out of the 32 amino acids are changed in comparison to N01. N04, the N-Cap of the DARPin domain of abicipar pegol ^27^.

## Results

### Choice of a minimal DARPin as model system

We chose a three repeat DARPin domain (denoted as N1C in the following) consisting of one full-consensus internal repeat (IR) flanked by an N- and C-Cap as a model DARPin to screen for improved thermostability (see **Table 1** for an overview of the different DARPin domains used, as well as **Table S1** for their respective sequences). The choice of this DARPin model has a three-fold motivation. First, it has the minimal DARPin architecture consisting of only three repeats with two repeat interfaces, one between the N-Cap and the IR and one between the IR and the C-Cap. Second, using a consensus IR that represents an “average structure” of all the natural ankyrin repeats should eliminate interferences originating from amino acids at randomized positions or possible framework mutations, that may only be present in particular sequences ^18^. Third, since DARPins get more stable with increasing number of IRs, we choose to have only one internal repeat and thus a low starting thermostability ^28^ such that stability improvements are readily observable.

**Table 1.** Description of DARPin domains used in the present study with varying N- and C-Caps as detailed in the text. For a comprehensive list of amino acid sequences of all domain variants used see **Table S1**.

| Domain name | Description |
| --- | --- |
| N1C | DARPin domain with one full consensus IR corresponding to NI1C of Interlandi et al. (2008)<sup>24</sup> & Wetzel et al. (2008)<sup>28</sup>. |
| aHER2 | DARPin domain based on an anti-HER2 DARPin domain corresponding to H10-2-G3 (“G3”; Zahnd et al. (2007)<sup>20</sup>). |
| aVEGF | DARPin domain based on an anti-VEGF-A DARPin domain of the protein moiety of abicipar pegol<sup>27</sup>. |
| aHSA | DARPin domain based on the anti-HSA DARPin domain of ensovibep<sup>10</sup>. It possesses an improved C-Cap similar to the mut5 C-Cap described by Interlandi et al. (2008)<sup>24</sup>. |

### Importance of the N-Cap position 17 for the overall thermostability of DARPins

By visual analysis of the DARPin structure, we identified four residues within the N-Cap to be of potential importance for the repeat stacking and thereby also for the overall domain stability. These residues are either at the edge (Leu4, Gly5, Asp17) of the N-Cap or buried (Met24) at the interface between N-Cap and the adjacent IR (**Figure 3**). As WO’655 already demonstrated that a Met24Leu mutation strongly improves the thermostability of DARPins, we focused our analysis on Leu4, Gly5 and Asp17 on an N1C background comprising the N02 N-Cap to find out if it is possible to further improve the most stable N-Cap known to date. Alanine scanning of residues 4 and 5 showed no improvement in thermostability, with Leu4Ala lowering the *T*_m_ value from 74.5°C to 64.7°C and Gly5Ala leaving the melting temperature unaltered at 74.3°C. However, the Asp17Ala mutation showed a strong improvement of the *T*_m_ value from 74.5°C to 82.4°C (**Figure 4A**). Consequently, we screened alternative amino acids at the N-Cap position 17 of N1C (**Table 2** and **Figure 4B**). All amino acid substitutions tested resulted in a *T*_m_ increase, with the highest *T*_m_ increases being measured for the Asp17Val, Asp17Ile and Asp17Leu variants (*i*.*e*. from 74.5°C to 85.1°C, 84.8°C and 84.6°C, respectively). Overall, changing Asp17 in N1C to Val, Leu, Ile, Met, Ala or Thr led to an increase of the respective *T*_m_ values between 8°C to 10°C. These results show that Asp is an exceptionally unfavorable amino acid at position 17 and that there are many alternative residues resulting in a strong thermostability gain. Of these alternative residues, Asp17Leu provided one of the largest improvements, which we investigated further.

**Table 2.** *T* _m_ values of N1C variants having various amino acids at position 17.

| Name | Position 17 | $T_m$ [°C] |
| --- | --- | --- |
| N1C_v01 | D | 74.5 |
| N1C_v04 | A | 82.4 |
| N1C_v05 | L | 84.6 |
| N1C_v06 | V | 85.1 |
| N1C_v07 | M | 83.8 |
| N1C_v08 | I | 84.8 |
| N1C_v09 | T | 82.3 |
| N1C_v10 | S | 79.3 |
| N1C_v11 | N | 75.2 |
| N1C_v12 | Q | 77.4 |
| N1C_v13 | K | 77.9 |
| N1C_v14 | R | 78.3 |
| N1C_v15 | E | 79.2 |

**Figure 3.**
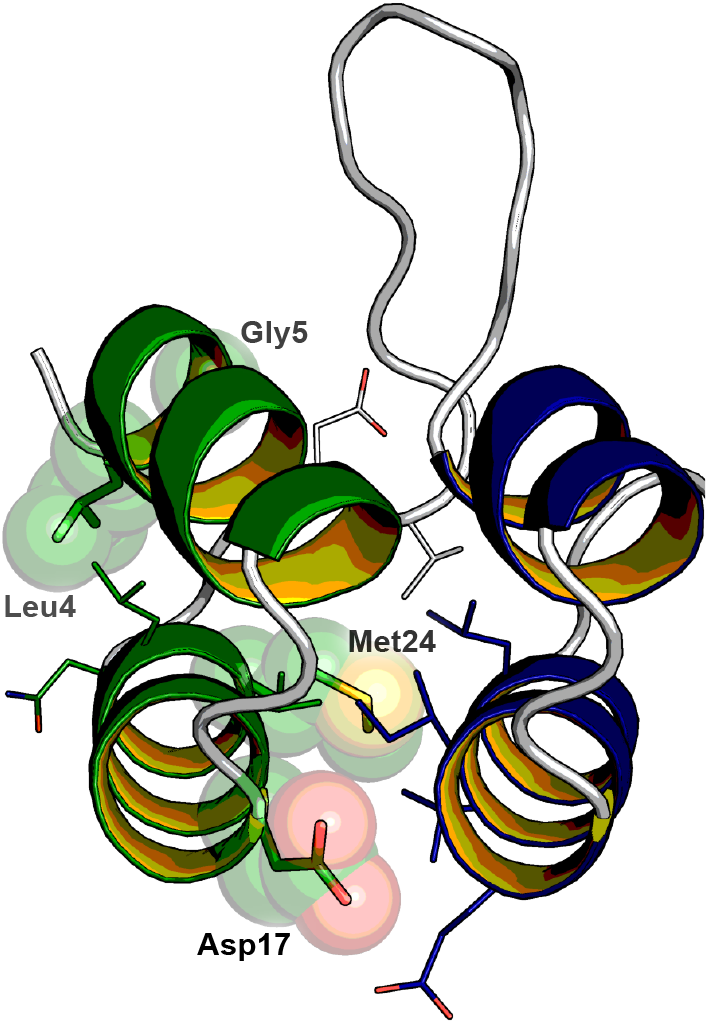
Ribbon diagram of an N01 N-Cap (green) and first IR (blue) of a conventional DARPin (PDB ID: 2XEE ^25^). Side chains of N-Cap residues Leu4, Gly5, Asp17 and Met24 are displayed as spheres and sticks and surrounding side chains are displayed as lines. This figure was created with open-source PyMOL ^26^.

**Figure 4.**
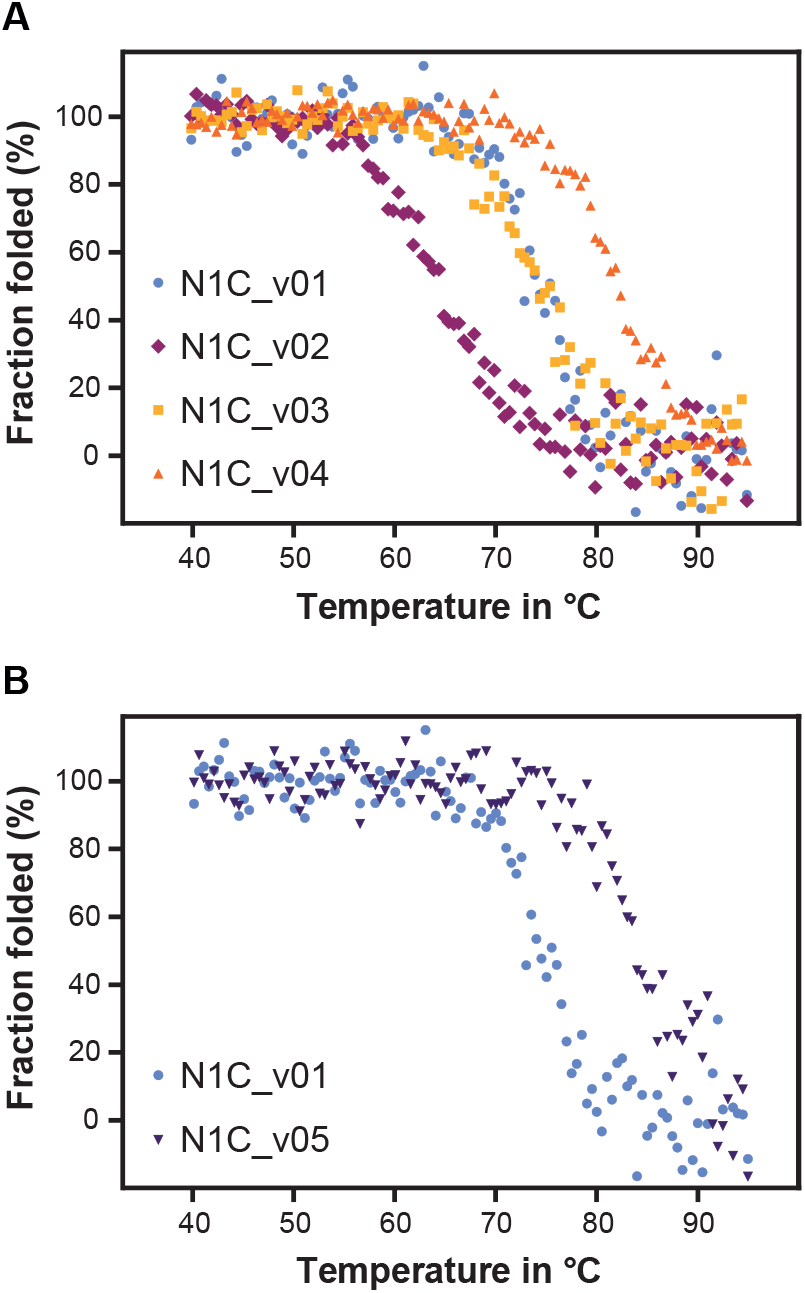
Thermal unfolding of DARPin domains followed by circular dichroism (CD) spectroscopy between 40°C and 95°C; all variants were measured at a concentration of 10 µM in PBS. **A)** N1C_v01 to N1C_v04: The measured *T* _m_ values were 74.5°C for the control N1C (N1C_v01), 64.7°C for the N-Cap Leu4Ala mutant (N1C_v02), 74.3°C for the N-Cap Gly4Ala mutant (N1C_v03) and 82.4°C for the N-Cap Asp17Ala mutant (N1C_v04). **B)** The Asp17Leu mutant (N1C_v05) has a *T* _m_ of 84.6°C and is strongly stabilized compared to N1C_v01 containing Asp17.

### The increased thermostability of the Asp17Leu N-Cap mutation is independent of the N- and C-Cap background

To test if the improvements derived from mutating the N- Cap Asp17 are generic and independent of the N02 background, we transferred the Asp17Leu mutation onto the original N01 N-Cap and the N03 N-Cap, that differs in nine amino acids from N02. The Asp17Leu mutation improved the thermostability of N1C also in the N01 and N03 backgrounds by more than 13°C (**Table 3**). Further, we were interested in whether our observed stability improvement based on the N-Cap and the stability improvement based on the mut5 C-Cap ^24^ would be additive. We first found that replacing the wt C-Cap in N1C with a mut5 C-Cap results in a *T*_m_ increase of about 13°C or 9°C in an N01 or N02 background, respectively (**Table 4**), thus confirming the benefits of the mut5 C-Cap ^24^. Additionally, the combination of the N02 N-Cap with the mut5 C-Cap in N1C_v22 proved that the individual improvements of each cap are additive, and raised the *T*_m_ value to 84.2°C in phosphate buffered saline (PBS), *i*.*e*. by about 22°C. With the additional substitution of Asp17Leu in N1C_v23 (N02, mut5 background) we did not observe any unfolding transition up to 95°C when we measured the thermal unfolding of this molecule in PBS. Therefore, we repeated the measurements for N1C_v22 and N1C_v23 in a buffer containing 2 M GdmCl and obtained corresponding *T*_m_ values of 67.3°C and 79.3°C, respectively (**Table 4**). Thus, the already very thermostable N1C_v22, comprising N02 and the mut5 C-Cap, could be further stabilized by adding the Asp17Leu mutation to its N-Cap resulting in a *T*_m_ gain of about 12°C in 2 M GdmCl. Overall, the Asp17Leu mutation adds about 9°C to 14°C to the *T*_m_ value of N1C independent of its concrete N- and/or C-Cap, indicating that this is a general improvement for DARPin domains.

**Table 3.** *T* _m_ values of N1C variants having either Asp or Leu at position 17 of the N-Cap in N01, N02 or N03 backgrounds.

| Name | N-Cap | C-Cap | $T_m$ [°C] |
| --- | --- | --- | --- |
| N1C_v01 | N02 | wt | 74.5 |
| N1C_v05 | N02_D17L | wt | 84.6 |
| N1C_v16 | N01 | wt | 62.1 |
| N1C_v17 | N01_D17L | wt | 75.2 |
| N1C_v19 | N03 | wt | 68.6 |
| N1C_v20 | N03_D17L | wt | 82.8 |

**Table 4.** *T* _m_ values of N1C variants having either Asp or Leu at position 17 of the N-Cap in wt and mut5 C-Cap backgrounds.

| Name | N-Cap | C-Cap | $T_m$ [°C] | $T_m^*$ [°C] |
| --- | --- | --- | --- | --- |
| N1C_v16 | N01 | wt | 62.1 | n.d. |
| N1C_v25 | N01 | mut5 | 74.6 | n.d. |
| N1C_v01 | N02 | wt | 74.5 | n.d. |
| N1C_v22 | N02 | mut5 | 84.2 | 67.3 |
| N1C_v23 | N02_D17L | mut5 | n.d. | 79.3 |

\*Indicates $T_m$ measurements in 2 M GdmCl.

### MD simulations suggest reduced flexibility of N1C through the Asp17Leu N-Cap mutation

We performed MD simulations to investigate the structural implications for N1C variants having either Asp or Leu at position 17 of their N-Caps. Starting from the X-ray diffraction structure of ankyrin repeat proteins of E3_5, NI1C-mut4 and NI3C-mut5 (PDB ID: 1MJ0^29^, 2XEN ^25^ and 2XEE ^25^, respectively) we prepared six different homology models, each time comparing Asp17-with Leu17-constructs in the N01-background (N1C_v16 and N1C_v17, respectively), the N02-background (N1C_v01 and N1C_v05, respectively), and the N02-background combined with the mut5 C-Cap (N1C_v22 and N1C_v23, respectively) (see **Table S1**). Starting from the homology models, three independent simulations were carried out for each system, two at 350 K and one at 400 K, for a total sampling of 1.8 µs. Three conclusions can be drawn from the MD simulations (**Figure 5** and **Table 5**). First, the substitution of Asp at position 17 with Leu leads in all instances to improved interaction energies with the surrounding (**Table 5**). Second, the analysis focused on the protein flexibility at high temperatures (400 K), revealed that the systems containing Leu at position 17 systematically show lower fluctuations than their Asp17 counterparts (**Figure 5**). The reduced flexibility of the Leu17 mutants as compared to Asp17 is in line with the increased thermostability of Asp17Leu DARPins observed in CD. Third, when examining the regions of reduced fluctuation, we found that the Asp17Leu mutation reduces fluctuations in the N01-background (N1C_v16 vs. N1C_v17) across the entire N-Cap and in one of the most flexible parts of a DARPin domain spanning from the end of the N-Cap (GADVNA motif, see (**Figure 2**) to the *β*-turn at the beginning of the internal repeat. In the N02-background (N1C_v01 vs. N1C_v05), reduced fluctuations through the Asp17Leu mutation are restricted to the direct vicinity of position 17 and around position 32, which is in line with the fact that N02 carries a stabilizing Met24Leu mutation. In the background comprising N02 and the mut5 C-Cap (N1C_v22 vs. N1C_v23) reduced fluctuations through the Asp17Leu mutation are only present in the direct vicinity of position 17. No fluctuation differences are observed around N-Cap position 32, which may be explained by the stabilizing effects introduced by the improved mut5 C-Cap that is structurally adjacent to this region. These findings further support our hypothesis that position 17 is an Achilles heel with regard to the N-Cap and thus the overall DARPin domain thermostability.

**Table 5.** Interaction energies between residue 17 and the rest of the simulation system (solvent and DARPin). The analysis was performed on the last 300 ns of each run at T = 350 K.

| Name | Coulombic [kcal/mol] | | van-der-Waals [kcal/mol] | | Total energy [kcal/mol] | | $T_m$ [°C] |
| --- | --- | --- | --- | --- | --- | --- | --- |
|  | Run 1 | Run 2 | Run 1 | Run 2 | Run 1 | Run 2 |  |
| N1C_v16 | -105.8 | -104.8 | -8.4 | -8.2 | -114.2 | -113.0 | 62.1 |
| N1C_v17 | -109.1 | -109.5 | -10.5 | -10.4 | -119.6 | -119.9 | 75.2 |
| N1C_v01 | -107.5 | -106.2 | -8.2 | -8.3 | -115.7 | -114.5 | 74.5 |
| N1C_v05 | -109.3 | -109.2 | -10.2 | -10.3 | -119.5 | -119.5 | 84.6 |
| N1C_v22 | -106.7 | -105.5 | -8.4 | -8.2 | -115.1 | -113.8 | 67.3* |
| N1C_v23 | -109.4 | -109.5 | -10.3 | -10.3 | -119.7 | -119.8 | 79.3* |

\*Indicates $T_m$ measurements in 2 M GdmCl.

**Figure 5.**
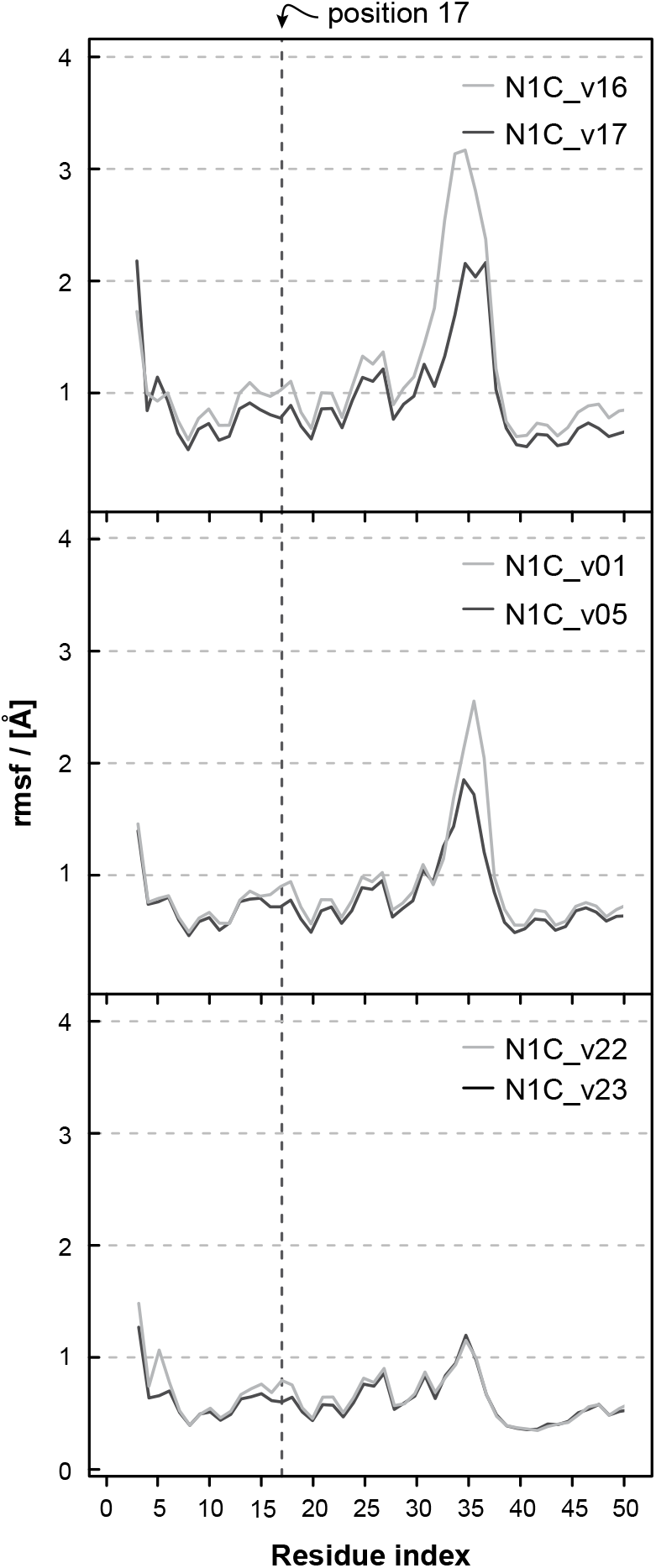
Root mean square fluctuations (RMSF) profile of the N-Cap and interface with the first internal repeat. Compared are the Asp17 (light grey) with the Leu17 (dark grey) constructs in the N01-background (top), the N02-background (middle), and the N02-background combined with the mut5 C-Cap (bottom). The last 300 ns of the 600-ns runs at 400 K were used to calculate the fluctuations of the Cα atoms. The stability of the folded structure of the Leu17-constructs is higher than the Asp17- constructs as the former show lower C*α* RMSF.

### The Asp17Leu N-Cap mutation improves the stability of clinically validated DARPins

To test whether the observed thermostability gain derived from the N-Cap Asp17Leu is independent on the composition of randomized positions in the DARPin paratope (as mainly present in the IRs), and thus transferable to any (non-consensus) DARPin, we tested this mutation on binders selected against human epidermal growth factor receptor 2 (HER2) ^20^, vascular endothelial growth factor A (VEGF-A) ^27^ and HSA ^30^ (**Table 6**). The selected DARPin domains are denoted aHER2, aVEGF (which carries a Gly5Asp framework mutation in its N-Cap; denoted as N04 in (**Figure 2**)) and aHSA, respectively (**Table 1**). Both aVEGF and aHSA are clinically validated DARPin domains as they are present in abicipar pegol and ensovibep, respectively. In all three trans fers, the Asp17Leu mutation increased the thermostability of the DARPin domains and added up to about 15°C in 2 M GdmCl *T*_m_ measurements (**Table 6**). Overall, the described significant gain in thermostability of the Asp17Leu mutation proved to be generic and is transferable to different N- and C-Cap backgrounds and different library members selected for high affinity, including clinically validated DARPin domains.

**Table 6.** *T* _m_ values of DARPin variants specifically binding to VEGF, HER2 and HSA.

| Name | N-Cap | $T_m$ [°C]<br>(in 2 M GdmCl) |
| --- | --- | --- |
| aVEGF_v01 | N04 | Unfolded at RT* |
| aVEGF_v02 | N04_D17L | 45.1 |
| aHER2_v01 | N01 | 39.6 |
| aHER2_v02 | N01_D17L | 55.7 |
| aHSA_v01 | N02 | 76.0 |
| aHSA_v01 | N02_D17L | 84.1 |

\*room temperature

## Discussion

The results presented in this paper identify position 17 of the DARPin N-Cap as a key position influencing thermostability of DARPins. The importance of the capping repeats as a pre-requisite for the high stability and robustness of the DARPin-fold has been known since the first description of this antibody mimetic ^28,29^. While the C-Cap has been thoroughly analyzed for residues providing room for improvement previously ^24^, an analogous dissection of the N-Cap in the literature, with exception of WO’655, has been missing.

### The N-Cap contributes significantly to the thermostability of DARPins

The contribution of the capping repeats to the overall stability of the DARPin domain is central ^28^. The architecture of solenoid repeat protein domains does not rely on long-range interactions (distant in sequence), which is a fundamental difference to globular proteins. In repeat proteins, stabilizing and structure-determining interactions are formed within a repeat and between neighboring repeats. In contrast to IRs, capping repeats have only one stabilizing neighboring repeat (**Figure 1**). Both natural ^31^ and designed ankyrin repeat proteins ^28^ have been shown to unfold in a sequential manner starting with the capping repeats. An unfolded cap leads to an internal repeat losing a stabilizing neighbor resulting in destabilization of the whole repeat domain. Furthermore, the original N- and C-Caps of DARPins correspond mainly to the natural caps of hGABP_beta1^2^; and thus there may be room for improvements. Indeed, improving the original capping repeats can increase the overall stability of a DARPin domain as it was shown for the N-Cap (WO’655) and the C-Cap ^24^. The improved N02 N-Cap of WO’655 comprises a Met24Leu mutation when compared to N01 (**Table 2**), thereby improving the thermostability of DARPin domains. Since Met possesses a larger side chain than Leu it might not optimally fit into the confined space at the interface between the N-Cap and the adjacent internal repeat. This mutation has the additional advantage that it eliminates an oxidation hotspot from the DARPin domain; unwanted oxidation of Met and its negative impact on the bioactivity of biologics is well documented ^32^.

We now performed a detailed analysis of the N-Cap, searching for additional mutations that may improve the overall DARPin domain stability. Following *in silico* analyses, we used equilibrium thermal denaturation experiments of DARPins harboring different N-Cap mutations. Intriguingly, our studies show that position 17 of the N-Cap is an important Achilles heel of a DARPin domain and that the original Asp at position 17 of the N-Cap is detrimental to overall DARPin thermostability. We showed that the negative effect on thermostability of Asp17 can be rescued by mutating it to Leu, Val, Ile, Ala, Met or Thr, leading to a profound improvement in *T*_m_ values by about 8°C to 16°C depending on the individual construct tested. Importantly, the *T*_m_ value of an N1C variant comprising the N02 N-Cap (instead of N01) was also increased by about 9°C when introducing the Asp17Leu mutation. Thus, the beneficial effect of this mutation synergizes with the *T*_m_ improvement of about 7°C previously reported for N02 (WO’655). While Interlandi et al. reported that the C-Cap is a limiting factor for the thermostability of the original DARPin domain ^24^, our results demonstrate that this is also the case for the N-Cap.

### Asp17 replacements seem to improve the N-Cap/IR interface

The N-Cap position 17 is located at the beginning of helix 2 at the edge of the interface between the N-Cap and the adjacent IR (IR1) (**Figure 6A**). Throughout nature, negatively charged amino acids are found at the N-terminal turn of helices as they support helix formation during protein folding ^33^, which rationalizes the placement of Asp at position 17. Known DARPin crystal structures show that the side chain of Asp17 can either face inwards and be buried in the interface between N-Cap and IR1, or face outwards into the surrounding solvent. If Asp17 is facing inwards, it is involved in van-der-Waals (VDW) contacts with IR1 (*i*.*e*. to the sidechains and backbones of IR1 Glu19 and IR1 Ile20), but is burdened with a desolvation penalty, which negatively impacts thermodynamic stability (**Figure 6A**). If facing outwards, Asp17 shields the hydrophobic interface from the solvent, but experiences repulsive forces to the equally negatively charged IR1 Glu19 (**Figure 6B**).

**Figure 6.**
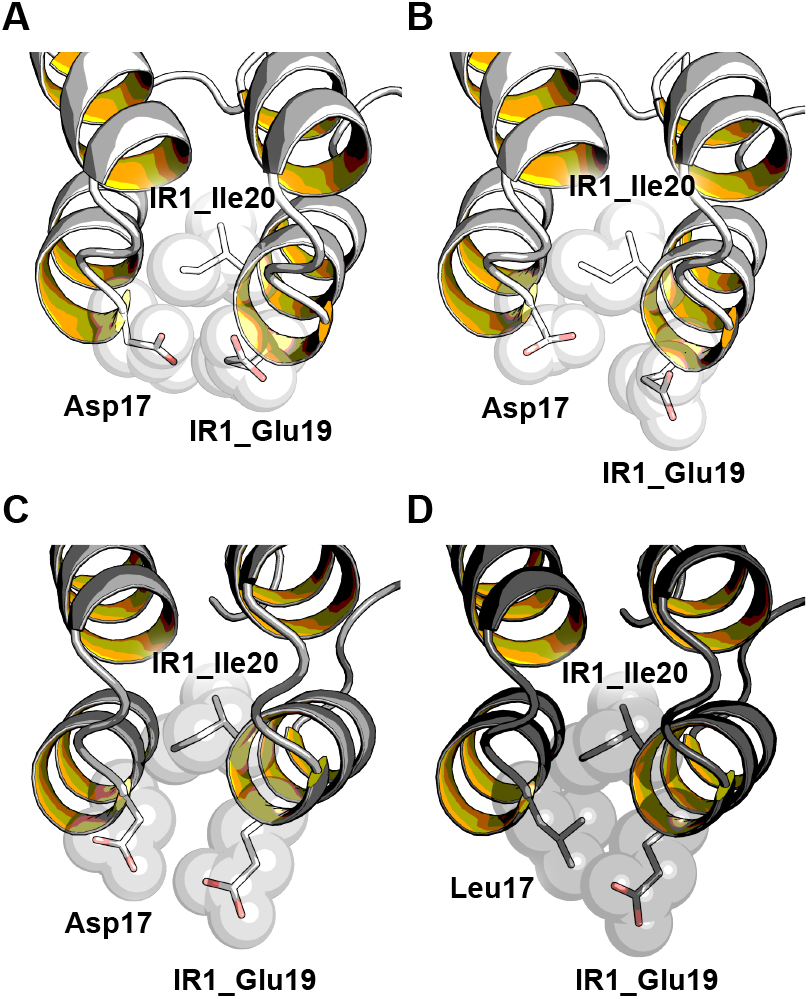
Side chain orientation and interaction of Asp17 and Leu17 with surrounding residues. **A)** Inward and **B)** outward facing Asp17 present in 2V4H ^34^. Average structures of the systems equilibrated at 300 K with **C)** outward facing Asp17 and **D)** inward facing Leu17. The N-Cap position 17 and internal repeat positions IR1_Glu19 and IR1_Ile20 are displayed as spheres and sticks. This figure was created with open-source PyMOL ^26^.

Analyses of equilibrated, energy-minimized average MD structures at 300 K and associated interactions between the N-Cap residue 17 and its surrounding at 350 K show that although in our starting model Asp17 faced inwards, during the course of the simulation Asp17 (**Figure 6C** and **Table 5**) is facing outwards as described above avoiding the desolvation penalty, but experiencing electrostatic repulsive forces to the IR1 Glu19. Importantly, the MD simulations show that Leu at position 17 faces inwards and that the Asp17Leu mutation improves the Coulombic and the VDW interactions by about 3 kcal/mol and 2 kcal/mol, respectively (**Figure 6D** and **Table 5**). The improved interactions of Leu17 are consistent with the surprisingly strong increase in thermostability observed by CD measurements.

### The Asp17Leu N-Cap mutation is universally applicable to DARPins

Importantly, the observed stability improvement of about 8°C to 16°C (depending on the concrete context) that was caused by the N-Cap Asp17Leu mutation, proved to be generically applicable to various sequence backgrounds. First, as observed by CD measurements and supported through MD simulations, stabilization could be consistently achieved on a diverse set of N-Cap backgrounds grafted on the model DARPin N1C. The tested N-Cap backgrounds include the original N01, N02 of the two HSA-binding domains of ensovibep, N03 where 12 out of the 32 amino acids are changed in comparison to N01, and N04, present in the VEGF binding domain of abicipar pegol (**Figure 2**). Second, the Asp17Leu mutation was also beneficial in the context of the improved mut5 C-Cap. Third, Asp17Leu gave a constant improvement when transferred to three different DARPin binders directed against HER2^20^, VEGF ^27^, or HSA ^30^, indicating that the observed profound improvement is independent of the randomized positions forming the DARPin paratope. This general transferability of the Asp17Leu mutation may be due to the fact that all DARPin domains are based on a quasi-identical framework embedding the randomized positions ^29^. The underlying reason for this similarity of the framework is the consensus design of DARPins, resulting in the stacking of self-compatible IRs ^1,2,18^. However, even the investigated DARPin domains aVEGF and aHER2 that possess framework mutations, which are key for their low picomolar binding affinities, still profit from the Asp17Leu mutation. This indicates that this single amino acid change may also be beneficial for DARPin domains comprising framework mutations; in particular, when these mutations have a negative effect on the domain stability.

Analyses of known X-ray structures of DARPins in complex with their respective targets have shown that N- and C-Caps contribute to the DARPin paratope in approximately 35% of cases ^35^. Position 17 is located outside of these known paratope regions and our MD based structural analyses indicate that the overall alteration of the domain structure through the Asp17Leu mutation is marginal. Thus, we do not expect that the N-Cap Asp17Leu change has a significant impact on the target binding of corresponding DARPin domains. Incorporation of the Asp17Leu mutation into existing DARPin binders could increase their thermostability without significantly impacting their target binding properties.

### Quantitative dimension and additive nature of the Asp17Leu improvement

The benefit obtained by this novel N-Cap (*i*.*e*., a *T*_m_ increase of 8°C to 16°C for a DARPin domain) is similar in dimension to that described for the C-Cap by Interlandi et al. ^24^. Most importantly, the N-Cap and C-Cap stability gains function in an additive manner, thereby yielding a *T*_m_ increase of approximately 20°C when combined. Similarly, the stability improvement of the novel N-Cap mutation Asp17Leu is additive to the stability gain obtained through the known N-Cap mutation Met24Leu (WO’655). When compared to the original DARPins ^1,2^, in sum, the two improvements at the N-Cap (each increasing the *T*_m_ value by approximately 10°C) and the improvement at the C-Cap can thereby yield a total *T*_m_ increase of approximately 30°C for a DARPin domain. Thus, our discovered Asp17Leu improvement adds substantially to other capping repeat improvements described previously. We see the modular, solenoid structure of DARPins as the origin for making the observed modular addition of thermostability improvements possible. DARPins are assembled through stacking of repeat units that are structurally “self-organized”, yet connected. They are stabilized by interactions with their folded immediate neighboring repeats and thus each repeat individually contributes to the overall domain stability ^28^.

### Ease of engineering of DARPins and impact on their versatility

DARPins unify a plethora of key characteristics beneficial for drug development from discovery to preclinical and clinical development, and manufacturing. For example, their low molecular weight, high solubility, high expression level, picomolar affinities, multispecific potential, and especially the high stability offer drug developers access to huge versatility ^9^. With their simple modular structure, multispecific DARPins ^16^ can be easily constructed with a single polypeptide chain and different molecular formats, as the binding domains are freely combinable. Their high solubility and high expression level awards them with unparalleled high-throughput capabilities and their excellent biophysical properties are the origin for low attrition rates. Their high thermostability, in particular, positively influences low aggregation and associated immunogenicity risks, allows for low-cost manufacturing and makes them very amenable to protein engineering to generate molecules with novel modes of action, such as receptor fine-tuning ^22^. With these characteristics DARPins complement existing therapeutic antibodies and will also expand the scope of innovative drugs via the multitude of advanced formats and applications that can be conceived and realized with this scaffold. All mentioned characteristics build on the stability of this versatile scaffold. Thus, an increased thermostability as presented in this study will have a positive impact on many levels.

### Significance for clinical DARPins

A recent example of drug development at unparalleled speed is the development of the multi-specific, highly efficacious anti-SARS-CoV-2 DARPin ensovibep that went from first selection of binders to entry into the clinic in less than nine months ^12^. Interestingly, ensovibep contains the Asp17Leu mutation described here in three of its five DARPin domains ^10^. This not only gives clinical validation to the Asp17Leu mutation, but also underscores the importance and reach of the findings outlined here. The high thermostability (*T*_m_ ⩾90°C) and absence of any tendency for aggregation (up to 85°C) reported for ensovibep ^11^ may thus be partly explained by the presence of the Asp17Leu mutation. These beneficial properties are especially remarkable as this COVID-19 antiviral drug is composed of five distinct DARPin domains with four different specificities on a single polypeptide chain. In our view, an immunoglobulin-based drug with similar properties would require extensive engineering to be generated, if feasible at all. Together, it does not surprise that Molecular Partners is speculating that ensovibep has the potential to bypass cold storage and that it provides a superior alternative to monoclonal antibody cocktails for global supply ^12^. Besides the three DARPin domains binding to three unique epitopes of the spike ectodomain of SARS-CoV-2, that all carry Asp17Leu in their N-Caps, ensovibep is additionally composed of two anti-HSA DARPins for serum half-life extension. These two domains are identical to aHSA of our study. As suggested by our findings, the overall stability of ensovibep might be even further improved by replacement of Asp17 with Leu, Val, Ile, Ala, Met or Thr in the N-Cap of the two anti-HSA DARPins.

Furthermore, aVEGF of our study corresponds to the ankyrin repeat domain of abicipar pegol that is a first generation DARPin drug to treat patients with neovascular age-related macular degeneration (AMD). Two randomized phase 3 clinical trials (CEDAR and SEQUOIA) demonstrated that quarterly applied abicipar pegol is noninferior to monthly applied ranibizumab, but showed a higher-level of intraocular inflammation (typically mild or moderate in severity) than ranibizumab ^36^. The authors of these phase 3 studies mentioned that the used abicipar pegol was further purified to reduce host cell proteins with the goal to minimize the incidence of intraocular inflammation. As indicated by our findings, the thermostability of the ankyrin repeat domain of abicipar pegol could be increased by about 30°C when combining our results with the previously described cap improvements. A correspondingly improved abicipar pegol would be amenable to more stringent purification processes that may result in much lower contaminations with host cell proteins.

## Conclusions

We have shown a *T*_m_ increase of various DARPin domains (up to 16°C) by replacing Asp17 in the N-Cap with Leu, Val, Ile, Ala, Met or Thr. Our results further provide evidence for the importance of the capping repeats for the robustness and reliability of this solenoid scaffold. Combining our N-Cap improvement with the cap enhancements described by Interlandi et al. ^24^ and WO’655, increases the melting temperature of original DARPin domains by about 30°C. Even though the initial DARPin design ^1,2^ is successfully used by the research community and is clinically validated (abicipar pegol), this significant improvement of the *T*_m_ indicates that the capping repeats of the original DARPins, which were based on hGABP_beta1, have high liabilities that can be eliminated by a few mutations in the capping repeats. Such thermodynamically stabilized antibody mimetics could pave the way for the future development of innovative drugs. First, increased thermostability of biologics is known to correlate with a reduced aggregation propensity ^37^. Thus, we anticipate an even lower immunogenicity risk of DARPins comprising the N-Caps described herein. Second, improved thermostability will make DARPins even more amenable to protein engineering. We believe that this is of high importance when the biological activity of a drug needs to be optimized, for example, to fine-tune the activation of receptors ^22^. Third, increased thermostability will translate into more efficient preclinical development and CMC processes, low-cost manufacturing and may even help to bypass cold storage of biologics. The recent development of the DARPin ensovibep (comprising the N-Cap Asp17Leu mutation) in less than nine months from idea conception to entry into the clinics underlines this huge potential ^11,12^. Along the same lines, we speculate that a high thermostability may facilitate the development of aerosol or dry powder inhaler formulations of DARPin drugs; something that may also be of interest for the pulmonary delivery of ensovibep for the treatment of COVID-19 patients. In conclusion, future drug development asks for more robust and very versatile biologics and we believe that DARPins would be the ideal scaffold for this.

## Methods

### Cloning

The DNA sequence encoding each ankyrin repeat domain was chemically synthesized and cloned into pQIq expression vectors (Qiagen, Germany) by Gibson Assembly ^38^.

### Protein expression and purification

The ankyrin repeat domains were expressed in *E. coli* XL1-Blue cells and purified using their His-tag by standard protocols. Briefly, 25 ml of stationary overnight cultures (LB, 1% glucose, 100 mg/l of ampicillin; 37°C) were used to inoculate 1 l cultures (same medium). At an absorbance of about 1 at 600 nm, the cultures were induced with 0.5 mM IPTG and incubated at 37°C for 4 h. The cultures were centrifuged and the resulting pellets were resuspended in 40 ml of TBS500 (50 mM Tris–HCl, 500 mM NaCl, pH 8) and sonicated. The lysate was recentrifuged, and glycerol (10% (v/v) final concentration) and imidazole (20 mM final concentration) were added to the resulting supernatant. The ankyrin repeat domains were purified over a Ni-nitrilotriacetic acid column (2.5 ml column volume) according to the manufacturer’s instructions (QIAgen, Germany). Up to 200 mg of highly soluble ankyrin repeat domains were purified from one liter of E. coli culture with a purity > 95% as estimated from SDS-PAGE.

### Circular Dichroism measurements

The circular dichroism (CD) signal of DARPin domains at 10 µM in PBS, pH 7.4 (PBS, 2 M GdmCl, pH 7.4, where indicated) were recorded at 222 nm in a Jasco J-810 instrument (Jasco, Japan) using a 1 mm pathlength cuvette. Samples were heated from 20°C to 95°C using a temperature ramp rate of 1°C per min, collecting data periodically at 0.5°C intervals. Melting temperature values were derived as described by Consalvi et al. ^39^. Importantly, all constructs assessed in CD ran as monomeric peaks in analytical size-exclusion chromatography (aSEC).

### Molecular Dynamics simulations

#### System preparation

Starting from the X-ray diffraction structure of ankyrin repeat proteins of E3_5, NI1C-mut4 and NI_3_C-mut5 (PDB ID: 1MJ0^29^, 2XEN ^25^ and 2XEE ^25^, respectively), homology modelling was used to construct atomic resolution models of six different repeat proteins, *i*.*e*., N1C_v16, N1C_v17, N1C_v01, N1C_v02, N1C_v22 and N1C_v23. The sequences of the proteins are shown in **Supplementary Table S1**. These models are used as the structural basis of this study. **Simulation protocol**. All simulations were carried out using the GROMACS 2018.6 simulation package ^40^ and the CHARMM36m force field ^41^. Three independent simulations with different initial random velocities were carried out for each repeat protein, two at 350 K and one at 400 K, cumulating 1.8 µs for each protein. To reproduce neutral pH conditions, standard protonation states were used for the ionizable side chains, while the N-terminus was positively charged, and the C-terminus negatively charged. Each protein was solvated in a cubic box (edge length of 6.9 nm) with TIP3P water molecules ^42^ to which 150 mM NaCl were added, including neutralizing counterions. Following steepest descent minimization, the simulation systems were first equilibrated under constant pressure for 5 ns, with position restraints applied on the heavy atoms, and subsequently under constant temperature (T = 300 K) in absence of restraints for 5 ns. For the production simulations, temperature and pressure were maintained constant at 350 K or 400 K and 1 atm, by using the modified Berendsen thermostat (0.1 ps coupling) ^43^ and barostat (2 ps coupling) ^44^. The short-range interactions were cut off beyond a distance of 1.2 nm and the potential smoothly decays to zero using the Verlet cutoff scheme. The Particle Mesh Ewald (PME) technique ^45^ with a cubic interpolation order, a real space cut-off of 1.2 nm and a grid spacing of 0.16 nm was employed to compute the long-range interactions. Bond lengths were constrained using a fourth order LINCS algorithm ^46^ with 2 iterations. In all simulations the time step was fixed to 4 fs, enabled through the use of virtual sites for all hydrogen atoms. Periodic boundary conditions were applied and the snapshots were saved every 50 ps. The elevated temperature is employed to enhance the sampling; the density of water is kept at the value corresponding to 300 K to perturb the free energy surface as little as possible ^47^.

## ACKNOWLEDGEMENTS

The authors would like to thank Dr. Lorenz Kallenbach, Dr. Nora Guidotti and Dr. Philipp Wild for critical reading and helpful comments during preparation and revision of the manuscript.

## DATA AND MATERIALS AVAILABILITY

The datasets generated and/or analyzed during the current study are available from the corresponding author upon reasonable request.

## AUTHOR CONTRIBUTIONS

JSchilling, CJ, IMI, JSchnabl, AC, PF: conceptualized and designed experiments; JSchilling, CJ, JSchnabl, OB, RSE, RT: performed laboratory experiments; IMI: performed molecular dynamics simulations; JSchilling, CJ, IMI, JSchnabl, RSE, AC, PF: drafted the manuscript. All authors reviewed the manuscript.

## Supplementary Tables

**Table S1.**
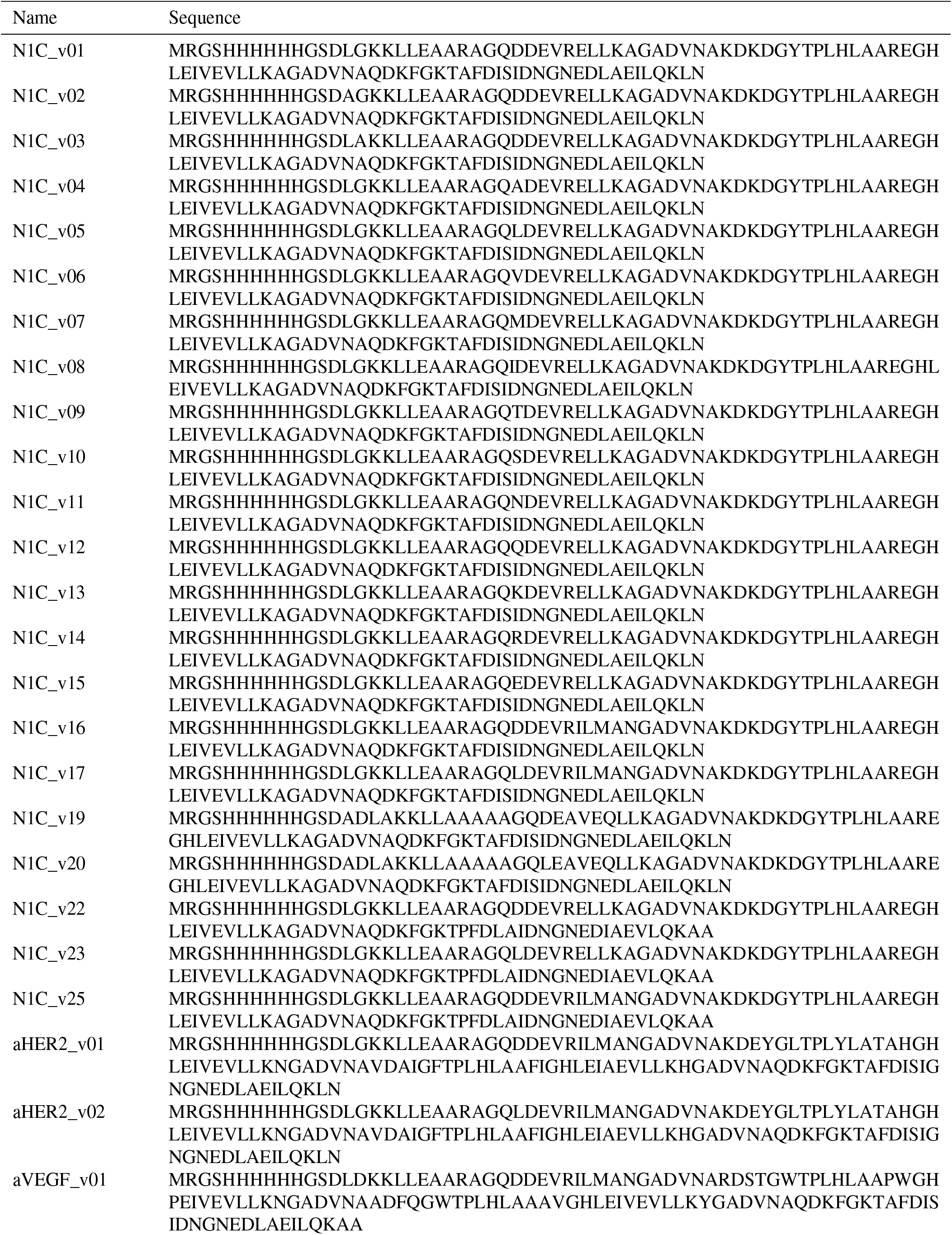
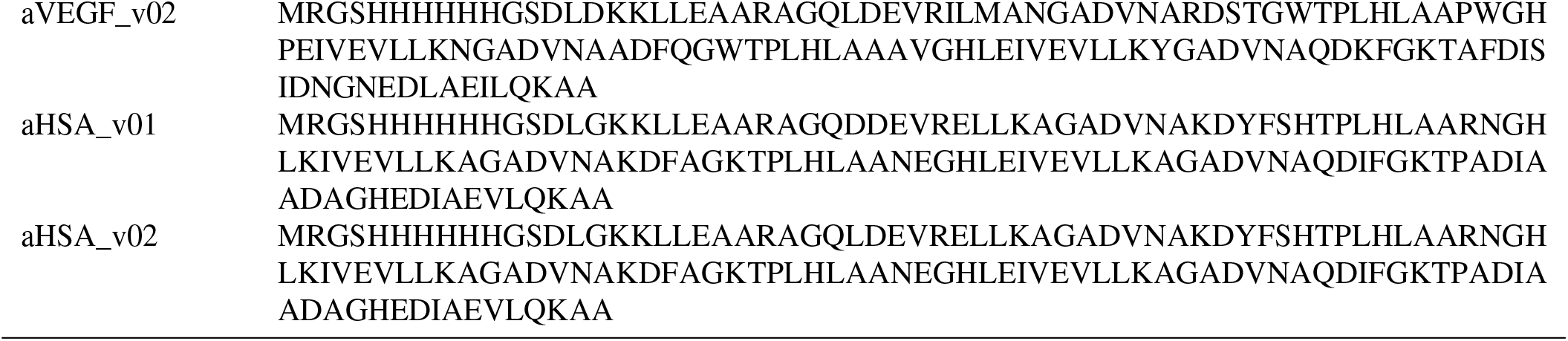
Protein sequences used in this study.

